# Multiannual environmental forcing shapes breeding phenology and success in a subantarctic seabird

**DOI:** 10.1101/2025.07.09.663721

**Authors:** Gaël Bardon, Téo Barracho, Joël M. Durant, Nicolas Lecomte, Yvon Le Maho, Nils Chr. Stenseth, Robin Cristofari, Céline Le Bohec

## Abstract

Climate-driven phenological mismatches threaten avian reproduction by disrupting food availability during critical breeding stages. In marine ecosystems, time lags between environmental changes and their effects on food webs are challenging to model, yet they can have a profound impact on top-predator reproduction. We disentangle how oceanic and climatic variability influence the breeding phenology and success of a keystone seabird of the Southern Ocean drawing on 24 years of data from 17,000 marked king penguins. We document an exceptional 19-day advancement in breeding phenology, alongside increased breeding success (44% in 2000, 62% in 2023). A sliding-window analysis reveals that sea temperature and primary production in key foraging zones predict both phenology and breeding success, with lags ranging from several weeks to two years. While king penguins appear to be keeping pace with current changes, their dependence on multi-year environmental conditions underscores the vulnerability of top predators to unpredictable, fast-changing, and more frequent extreme conditions.

## INTRODUCTION

The timing of life-history events like reproduction is central in optimising fitness, even more so under changing environmental conditions, as it effectively modulates the biotic and abiotic conditions and interactions encountered by both parents and offspring (McNamara et al., 2011; Miller-Rushing et al., 2010) However, the phenology of high trophic level species is often less responsive to changing conditions than that of low trophic level species such as primary producers (Both et al., 2009; Thackeray et al., 2010). This means that, while their prey might follow the changes in climate, top-level predators could be unable to keep pace by adjusting their breeding, growth cycles and distribution (Burthe et al., 2012), leading to a well-documented mismatch between peaks in resource availability and breeding phenology (Burgess et al., 2018; Durant et al., 2007; Kharouba et al., 2018).

At high latitudes, ecological processes such as thermal regimes, photosynthesis, and behaviour of visual organisms are strongly seasonally constrained by light (Ljungström et al., 2021). This seasonal light availability and temperature shape a narrower annual peak of productivity compared to tropical and subtropical zones (Boyce et al., 2017; Falkowski et al., 1998), leading to a high risk of trophic mismatch in the current context of rapid global change (Durant et al., 2019; Ljungström et al., 2021). Many high-latitude species are already in mismatch situations (e.g., Doiron et al., 2015; Kwon et al., 2019). Combined with the fact that polar environments are experiencing among the fastest warming on Earth (IPCC, 2022), this represents a critical threat to these ecosystems.

In marine food webs, extreme abiotic forcing and high capacity for the displacement of primary producer blooms make decoupling of trophic levels prevalent (Durant et al., 2019; Ferreira et al., 2023), leading to bottom-up impacts on higher trophic levels (Asch et al., 2019; Yamaguchi et al., 2022). Seabirds in high latitudes, as top predators, are particularly sensitive to changes in environmental conditions because their breeding success depends on synchronising their limited seasonal reproductive window with prey availability (Burr et al., 2016; Descamps et al., 2019; Seyer et al., 2021). Yet, despite observed strong changes in abiotic forcing in polar oceans and phenological shifts in the trophic network (IPCC, 2022; Yamaguchi et al., 2022), many seabirds have not altered their breeding phenology (Keogan et al., 2018) or with little consistency with prey phenology (Burthe et al., 2012). In Arctic-breeding seabird populations heavily impacted by climate change, thick-billed murres *Uria lomvia* have advanced their laying dates in response to environmental shifts. However, this adjustment has lagged behind changes in prey phenology, leading to a mismatch with the peak of high-quality prey availability (Whelan et al., 2022). Rapid changes in the Southern Ocean (IPCC, 2022) may result in observable mismatch situations for seabirds breeding there. By examining the relationship between phenology, breeding success, and the impact of environmental changes, our aim is to gain a deeper understanding of the underlying mechanisms for avoiding mismatches and their limitations.

This study focuses on one of the main central-place foragers of the Southern Ocean, the King penguin *Aptenodytes patagonicus*. King penguins establish breeding colonies of a few hundred to several hundred thousand individuals on the ice-free coastal lands of the few sub-Antarctic islands (45°S to 55°S). Despite this restricted breeding latitudinal range and environmental conditions, the species displays notable plasticity in its phenology, behaviour, and life-history traits (Bost et al., 2013). In response to fluctuations in prey availability, king penguins can adjust key aspects of their foraging strategy, such as shifting their foraging latitude in relation to the Antarctic Polar Front (Bost et al., 2009, 2015), modifying trip duration (Weimerskirch et al., 1992), or increasing foraging effort (Brisson-Curadeau et al., 2024). They also show considerable individual phenological flexibility, with the ability to initiate successful breeding between November and February. However, the extent to which king penguins adjust their breeding phenology in response to environmental variability remains poorly understood, as well as the consequences of such adjustments for breeding outcome. Phenological plasticity likely depends on the ability to detect and respond to environmental cues—a capacity increasingly challenged by the rapid and complex environmental changes occurring in the Southern Ocean. These changes encompass both large-scale trends, such as the positive shift in the Southern Annular Mode (King et al., 2023), and localised, often contrasting, shifts in biogeochemical and climatic conditions (Fogt & Marshall, 2020; Henley et al., 2020), with cascading effects on trophic dynamics (Krumhardt et al., 2022; Thomalla et al., 2023). These rapid and complex environmental changes increase the risk of phenological mismatches in sub-Antarctic species. To unravel how oceanic and climatic conditions influence the breeding activities of king penguins, we used data from a 24-year-long electronic monitoring program in the subantarctic archipelago of Crozet in the Southern Ocean (Supp. Figure S1), home for 20 to 25% of the King penguins’ world population (Barbraud et al., 2020; Bost et al., 2013). After determining the annual breeding phenology and the annual breeding success of the study population, we examined which of a wide range of indices best explained the variability in the breeding timing and outcome, and over which time window.

## RESULTS AND DISCUSSION

### Phenological and breeding success trend

We observed a temporal trend in annual phenology towards an earlier onset of breeding. The breeding season shifted eight days earlier per decade over the 24-year study period (27 November in 2000 to 8 November in 2023; slope = 0.81 ± 0.23 days per year, p-value = 0.0016, Figure 1A). Over the same period, breeding success increased slowly at an annual rate of 0.76% (44.2% in 2000 to 61.7% in 2023; slope = 0.76 ± 0.31% per year, p-value = 0.0232, Figure 1B), which was strongly correlated with the median breeding date (slope = −0.97± 0.16 % per day, p-value << 0.001, Figure 1C). No temporal trend in breeding success was observed when the median breeding date was accounted for in the model (p-value = 0.885). However, substantial interannual variation remained (residual standard error = 7.4%), indicating that the median breeding date explains only part of the annual variation in breeding success.

**Figure 1.**
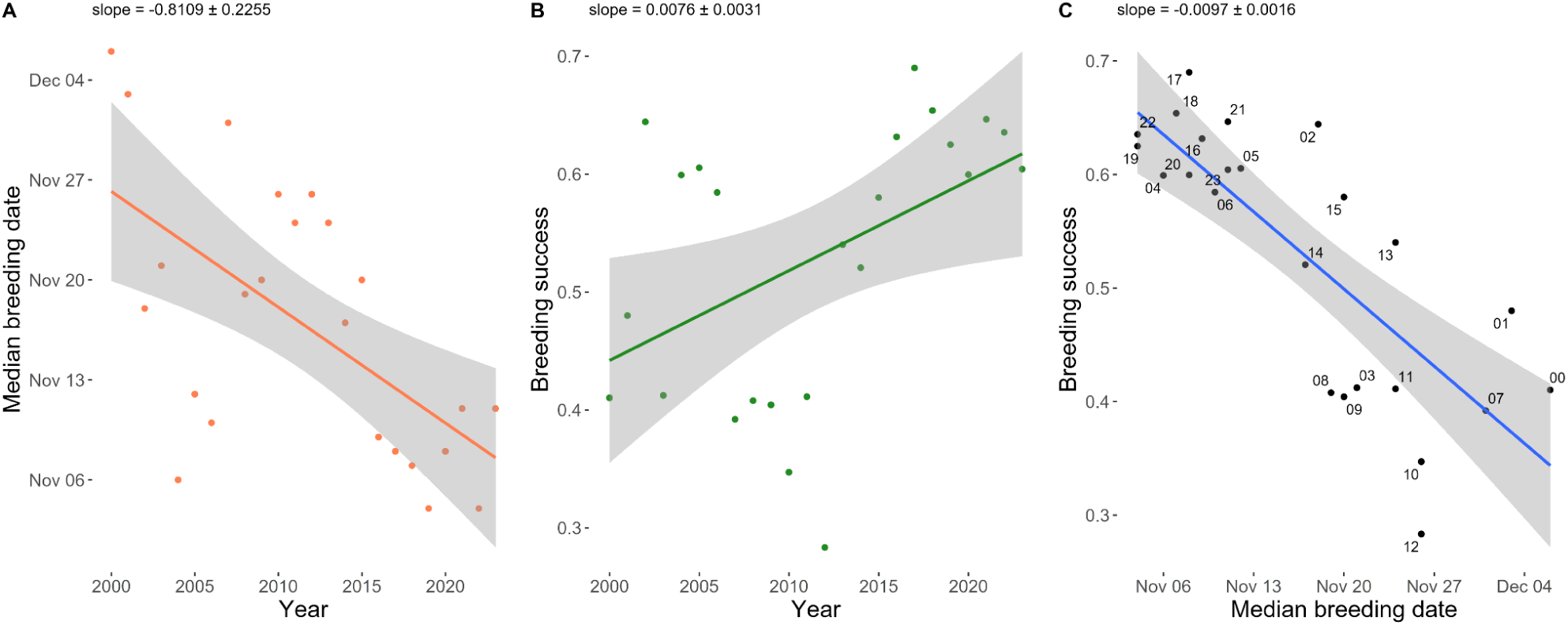
Temporal trends in king penguin breeding phenology and success of the Possession Island, Crozet archipelago, South Indian Ocean. (A) Annual median breeding dates, (B) annual breeding success, and (C) annual breeding success in relation to the median breeding date. Each point represents one year, and the blue line is the univariate linear regression predicted average. The grey shaded areas show 95% confidence intervals of the regression lines. Slope coefficients and their standard error are given for each regression.

This 19-day advancement in phenology over the past 24 years, corresponding to an 8.1-day decadal advance, is remarkable compared to other high-latitude species (Cohen et al., 2018). For example, the decadal advance in breeding initiation reached only 1.25 days (95% CI: 0.925, 1.375 days) for 73 bird species in Finland (Hällfors et al., 2020), on average 1.2 days (ranging from 0.3 to 2.3 days) for Swedish Lapland passerines (Ram et al., 2019), or 4 days for Arctic squirrels *Urocitellus parryii* (Chmura et al., 2023). To our knowledge, it is also the fastest shift observed in penguin species. The clutch initiation date of Adélie penguin *Pygoscelis adeliae* from the Antarctic Peninsula has remained stable over the last three decades despite major changes in environmental conditions (Cimino et al., 2023; Youngflesh et al., 2017). For royal penguins *Eudyptes schlegeli*, an average advance < 1 day per decade has been recorded over 35 years (Hindell et al., 2012).

King penguins appear to succeed in tracking favourable environmental conditions for reproduction. Our results indeed suggest that recent environmental changes may have been beneficial to the breeding success of the Crozet king penguin population, enabling earlier breeding by reducing constraints on the breeding season—albeit with a significant interannual variability in this trend. The magnitude of this effect is substantia: each day’s advance in breeding timing corresponds to approximately 1% increase in breeding success, with significant consequences in terms of population productivity.

Studies have shown that climate-driven advances in breeding timing can have contrasting effects on the reproductive success of migratory and sedentary bird species (Dunn et al., 2010). In some cases, earlier breeding may be detrimental: for example, black grouse *Lyrurus tetrix* in Finland have shifted their hatching dates in response to warmer springs, but early summer temperatures have not increased accordingly. This mismatch exposes chicks to colder conditions, leading to higher mortality rates and reduced reproductive output (Ludwig et al., 2006). Conversely, breeding success may remain unaffected by an advance in phenology, suggesting that some species are able to maintain synchrony with optimal environmental conditions during key breeding stages (Dyrcz & Czyż, 2018; McLean et al., 2022; Usui et al., 2017). However, many species may benefit from phenological advances —for instance, multibrooded species that can extend their breeding season (McDermott & DeGroote, 2016). This is further supported by a translocation experiment of pied flycatchers *Ficedula hypoleuca* to the north: translocated females, that bred earlier than the local females, exhibited a 2.5 times higher fitness, suggesting that the local population had not yet fully optimised its breeding phenology (Lamers et al., 2023). In our study, no evidence of trophic mismatch has been observed, suggesting that king penguins can synchronise the most sensitive periods of their reproductive cycle in terms of chick-provisioning with maximum prey availability. King penguins have a prolonged breeding cycle lasting 12 to 14 months, which includes a 3-month winter fasting period for chicks (Barrat, 1976). During this time, chicks are exposed to harsh environmental conditions, making it a phase of increased mortality. An advance in breeding phenology may allow chicks to attain better body condition before winter, a factor known to enhance overwinter survival (Brisson-Curadeau et al., 2023) and, consequently, breeding success.

Yet, with the ongoing fast and projected changes in Southern Ocean ecosystems, major shifts in the marine food web might in result in seasonal mismatches between predators and prey at multiple trophic levels (Yamaguchi et al., 2022), threatening penguins (Cimino et al., 2023), their prey, and the ecosystem as a whole (Durant et al., 2019).

### Environmental predictors of breeding phenology

Seasonally predictable environmental cues, in other words, phenological cues, that guide the annual breeding initiation of high-latitude top predators, have been largely unexplored. Identifying the variables that are strongly correlated with the breeding initiation date could therefore provide some initial clues as to the information used by the birds to initiate breeding. Thus, we investigated how environmental conditions (i) around king penguin breeding habitat (ii) in their main foraging areas, and (iii) at a global scale, impact the date of their annual breeding onset while accounting for short- and long-term lags (see tested variables and model selection in Supp. Table 1 and Supp. Table 2). For that purpose, we used a sliding window analysis (Bailey & Van De Pol, 2016; Van De Pol et al., 2016; see Methods). To capture a wide range of potential relationships between environmental conditions and breeding initiation, we tested both linear and quadratic effects. The inclusion of quadratic terms allowed us to assess the possibility of non-linear responses, particularly the existence of optimal environmental conditions. Such optima may reflect thresholds beyond which environmental changes become detrimental to prey abundance or accessibility.

Using the sliding window analysis combined with model selection, we identified two environmental variables associated with the timing of breeding initiation in king penguins: (a) Chlorophyll *a* concentration (Chl*a,* Figure 2B) in late winter (i.e. mid-August to late September preceding the breeding onset) with a linear effect, and (b) the Sea Surface Temperature (SST, Figure 2A) in autumn (i.e. the first two weeks of May preceding the breeding onset) with a quadratic effect, both in the Polar Front Area (i.e. the main summer foraging area for the species; see Supp. Figure 1). These two variables explained 82% of the temporal variance in breeding phenology, though model uncertainty increases when projecting beyond the observed range of environmental conditions. Also, none of these two variables taken in isolation showed a significant temporal trend over the study period (Supp. Figure 7). However, when accounting for these two environmental variables, the year coefficient is not significant in the final model (slope = −0.25 ± 0.14 days per year, p-value = 0.087), unlike in the year-only model (slope = 0.81 ± 0.23 days per year, p-value = 0.0016). This observation clearly indicates that a substantial fraction of the observed trend in breeding phenology is explained by the combination of these two variables.

**Figure 2.**
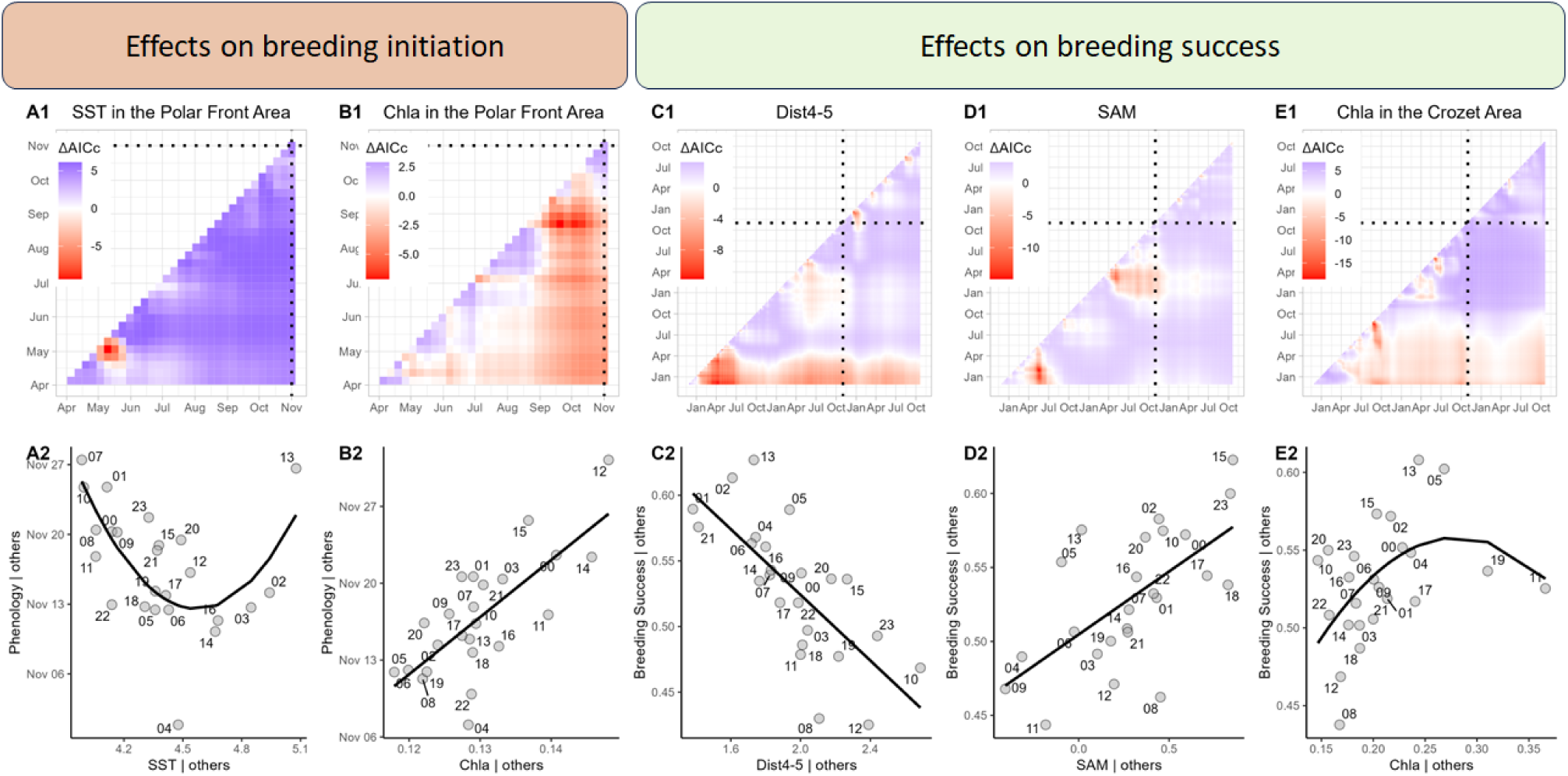
Environmental conditions affecting the king penguin breeding initiation date and breeding success. The first two columns (A and B) correspond to the variables selected to explain the breeding initiation date. The three columns on the right (C, D, E) correspond to the variables selected to explain the breeding success rate. The dashed lines correspond to the average breeding initiation date. A1 and B1 represent the ΔAICc between the reference model explaining breeding initiation date (Median breeding initiation date ∼ Year) and the same model including the mean value of the environmental variable over each time window. C1, D1, and E1 represent the ΔAICc between the reference model explaining breeding success (Breeding success ∼ Year + Median breeding date) and the same model including the mean value of the environmental variable over each time window. The time windows are given by the coordinate of each pixel: the window starts at the coordinate of the y-axis and ends at the coordinate of the x-axis. Bottom panels: Partial regression plots showing the isolated effect of each environmental variable on breeding timing/success. Points represent residuals after removing effects of all other model variables, with means added back for interpretability. These plots reveal the unique contribution of each predictor (see Methods for details). SAM: Southern Annular Mode Index; Chla: Chlorophyll a concentration; SST: Sea Surface Temperature; Dist4-5: Distance between 4°C and 5°C isotherms. The Crozet Area (43°S-47°S - 46°E-56°E) and the Polar Front Area (47°S-53°S - 49°E-55°E) are the areas used to calculate our environmental variables.

Lower Chl*a* and an optimal SST in the Polar Front Area were associated with an earlier onset of breeding. SST is a well-established driver of fish productivity across numerous species (see review in van der Sleen et al., 2022). The seasonal growth peak of several species of myctophidae, which are primary food sources for king penguins (Cherel et al., 1993), has been observed to occur in autumn (Saunders et al., 2020). The optimal SST (i.e. the minimum of the quadratic relationship at ∼4.5°C, see Figure 2A) identified in our study likely reflects oceanographic conditions near the Polar Front before winter, that enhance myctophid accessibility to foraging penguins, though the mechanistic link requires further investigation. The negative relationship between Chl*a* and myctophidae abundance may seem counterintuitive but reflects the complex trophic dynamics of the Southern Ocean. High surface chlorophyll concentrations can indicate either highly productive but unstable conditions or mismatched timing between phytoplankton blooms and zooplankton grazing (Loots et al., 2007). Myctophidae such as *Electrona antarctica* and *Protomyctophum tenisoni* - key prey species for king penguins in winter and spring - may be more abundant when Chl*a* levels are moderate, suggesting optimal rather than maximum primary productivity supports their populations (Woods et al., 2023). Additionally, high Chl*a* may occur in areas with poor physical oceanographic conditions for fish aggregation, such as areas with weak frontal structures.

As recently shown for black-legged kittiwakes *Rissa tridactyla* (Whelan et al., 2022), the phenology of king penguins is likely driven by bottom-up effects of environmental conditions rather than direct environmental cues. When conditions favour an increased abundance of their primary prey, king penguins may attain adequate body condition earlier in the spring, thereby allowing an earlier onset of moulting and breeding. However, surface ocean Ch*a* has already been identified as likely phenological cues for two auk species (Crossin et al., 2022), suggesting a mechanism that could potentially apply to king penguins as well. Nevertheless, it remains unclear what information this species could derive from Ch*a* during winter and how this might influence its reproduction.

### Environmental predictors of breeding success

Using the same sliding window analysis method, we found that, while breeding phenology explained a substantial part of the variance in breeding success (Figure 1), the remaining variance was mainly explained by three environmental parameters (Figure 2D-E-F) with a time lag of a year and a half to two years prior to the onset of the breeding season (see model selection in Supp. Table 3). Our selection model indicates (a) a linear effect of the distance between the isotherms 4°C and 5° (mid-December to mid-April, ∼2 years prior), (b) a linear effect of the Southern Annular Mode (SAM) index (February to May, ∼1.5 years prior), and (c) a quadratic effect of the mean Chlorophyll *a* concentration (Chl*a* in mg/m^3^) in the Crozet Area in late winter (mid-August to early September, ∼1 year prior). These three variables together with the phenology explained 94% of breeding success variance.

A short distance between the 4°C and 5°C isotherms (i.e. a strong latitudinal SST gradient) and a positive SAM index appear to benefit king penguin breeding success. Both the strong SST gradient and the positive SAM are associated with positive temperature anomalies in the subtropical zone and negative anomalies south of the Antarctic Polar Front (Fogt & Marshall, 2020), a pattern associated with increased primary productivity in Antarctic waters (Lovenduski & Gruber, 2005). King penguin breeding success showed a quadratic relationship with Chl*a* in the Crozet Area (Figure 2E), with highest success at intermediate levels (0.26 mg/m³) slightly over one year before the start of the breeding cycle, though the confidence interval around this optimum spans 0.20-0.32 mg/m³. Derived from satellite imagery, Chl*a* reflects only surface-layer phytoplankton density and does not linearly correlate with total primary productivity (Westberry et al., 2008). We thus suggest that Chl*a* optimum is linked to optimal conditions that favour the growth and abundance of available prey, and especially squids (Cherel & Weimerskirch, 1999), spawned near the breeding colony. King penguins undertake two types of foraging trips: long-distance trips to the Polar Front Area, located approximately 300 to 500 km south of the colony (Bost et al., 2015), and shorter, local trips around the colony, where prey distribution is typically more variable and occurs at greater depths (Jouventin et al., 1994). Increased prey availability in the vicinity of the colony may enhance foraging efficiency and reduce the duration of foraging trips, ultimately supporting greater breeding success.

The 1-2 year temporal lag between these three environmental conditions and breeding success likely reflects two complementary mechanisms. First, given that king penguins feed primarily on subadult and adult myctophids and squids (Bost et al., 2002; Cherel & Ridoux, 1992), environmental conditions during prey spawning and recruitment phases (1-2 years prior) determine the abundance of fish cohorts that become available during subsequent breeding seasons. Second, the king penguin’s extended breeding cycle (12-14 months) means that environmental effects on adult body condition and energy reserves can carry over to influence breeding decisions and performance in following seasons. These mechanisms may act synergistically, with favourable environmental conditions enhancing both prey availability and parental condition over multi-year timescales It has been reported that warm events (depicted by negative SOI) in any given year were linked to reduced king penguin breeding success (Le Bohec et al., 2008) and that positive anomaly of SST in the Polar Front Area could also negatively affect breeding success with penguins travelling further to reach the high productivity located at the oceanic fronts (Bost et al., 2009, 2015). Unlike these previous studies on the same penguin population, we found no relationship between breeding success and warm environmental conditions during the ongoing breeding season. This suggests that the abundance and/or quality of prey, driven by oceanographic conditions up to one to several years prior to a breeding season, may play a more decisive role than prey spatial distribution. Except in extreme years, such as 1997 when the Polar Front shifted too far south to support successful breeding (Bost et al., 2015), our findings underscore the capacity of king penguins to adjust their foraging strategies and breeding phenology in response to environmental variability.

To conclude, our study shows a drastic advance of the breeding season by two weeks over the last two decades in a key species of the Southern Ocean ecosystems usually considered as an indicator of environmental changes (Durant et al., 2009; Le Bohec et al., 2008; Parsons et al., 2008; Piatt & Sydeman, 2007). This result pinpoints profound and rapid changes trickling all the way to the top of the marine food chain, by increasing the breeding success of the king penguin populations of the Indian Ocean sector. King penguins from the Kerguelen archipelago, located 1,500 km east of Crozet, also appears to benefit from the current rise in temperature through a positive effect on prey abundance (Brisson-Curadeau et al., 2023). For now, king penguins appear to benefit from current climate regime in the Southern Ocean, alongside around 20% of benthic species (Griffiths et al., 2017) and several seabirds and tuna species whose core habitats are projected to expand (Hazen et al., 2013; Reisinger et al., 2022). Then, organisms seem to keep pace, matching their needs with available resources and responding to changing environmental conditions, but for how long? How much of this pace is temporary and context-dependent? The plasticity of populations and their ability to cope with future changes is still challenging to gauge (e.g., Lewin et al., 2024). Non-linear response of species to changes in their environment might be common. For example, increased shrub cover and density, favoured by climate change, may potentially be beneficial for some Arctic tundra-breeding bird species; but as shrub height increases further, however, a considerable number of these birds will likely find habitat increasingly unsuitable (S. J. Thompson et al., 2016). The quadratic effects found in our study provide an initial understanding of the Southern Ocean ecosystem’s limits for king penguins, indicating negative impacts from excessively warm temperatures (> 4.6°C in the Polar Front Area in May, Figure 2A) and too high chlorophyll concentrations (> 0.50 mg/m^3^ in the Crozet Area in the end of August, Figure 2E). Southern Ocean climate projections (e.g. 1-3°C warming, front displacement, increased SAM intensity; IPCC, 2022) suggest that environmental conditions may exceed the optimal ranges identified here within 2-3 decades. However, we hypothesize that the 1- to 2-year lag between environmental conditions and breeding responses observed in our study may obscure early signs of declining habitat suitability, delaying the detection of population-level impacts.

We have also highlighted that the initiation and success of the king penguin breeding cycle result from the integration of environmental conditions spanning a period of up to two years around their breeding and foraging grounds (Figure 3). Short-term conditions just prior to the breeding season might drive its onset by facilitating earlier, generally longer and more successful breeding under favourable foraging conditions. They may act as direct phenological cues informing penguins when to start breeding, which generally guarantees them to be synchronised with their prey’s phenology, as observed over the past 24 years. Notably, our findings contrast with those of Keogan et al. (2018), who reported that seabirds globally do not exhibit phenological shifts in response to sea temperature variability — a conclusion that may overlook the role of temporal lags in species’ responses to environmental change. Nevertheless, at current rates of change, king penguins may not be able to integrate all the information from two years of oceanographic data at the right pace, making the prediction of resource availability virtually impossible. Growing evidence supports the existence of such long-term environmental response lags in various species, including murres, kittiwakes, and cormorants in the North Pacific and Atlantic Oceans (Whelan et al., 2022; Zador et al., 2013). Our study extends these observations to the Southern Ocean and provides a clear illustration of the bottom-up trophic chain consequences of climate change, demonstrating how environmental variability at the base of the food web can propagate through trophic levels to influence top predators such as king penguins. With the more frequent occurrence of extreme and unprecedented events resulting from climate change (IPCC, 2022), we have yet to understand the consequences of species inertia in their response to fast-changing and more extreme weather patterns. So far, the 2-year lag in king penguin response in breeding output is still allowing reproductive benefits whether this translates into juvenile survival and ultimately population dynamics is unclear. Future work linking the length of lag responses to population trajectories and ecosystem stability is a welcome challenge for deciphering biodiversity changes with current and future climatic conditions.

**Figure 3.**
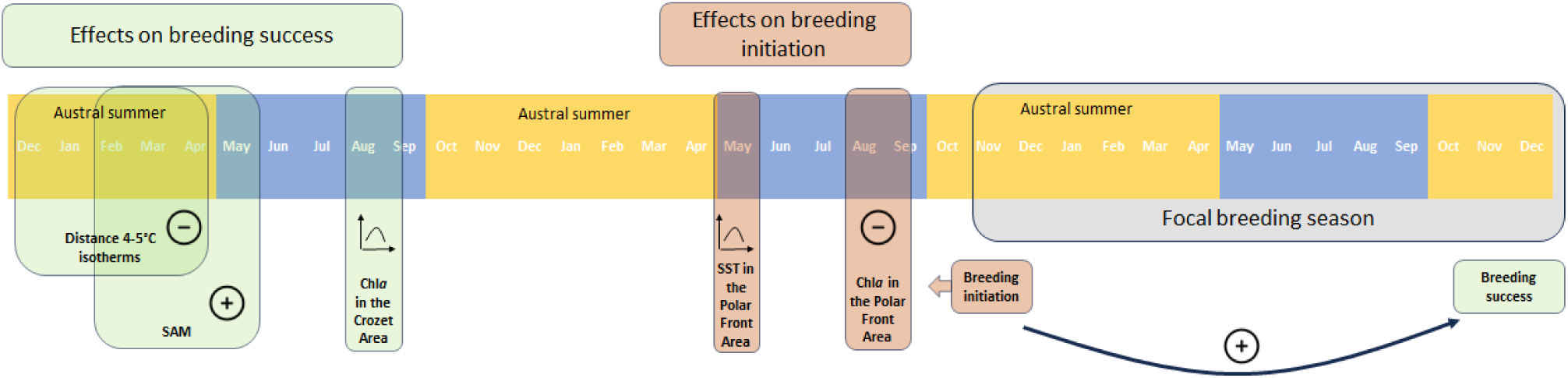
Overview of the relations found between environmental conditions, breeding phenology, and breeding success in king penguins. SAM: Southern Annular Mode Index; Chla: Chlorophyll a concentration; SST: Sea Surface Temperature. The Crozet Area (43°S-47°S - 46°E-56°E) and the Polar Front Area (47°S-53°S - 49°E-55°E) are the areas used to calculate our environmental variables. ‘Plus’ indicates that the variable has a positive effect either on the breeding success (in green) or on the breeding initiation (in red). In this figure, a positive effect on breeding initiation corresponds to an earlier onset of breeding initiation. The black arrow indicates a positive correlation found in the current study.

## MATERIALS AND METHODS

### Long-term monitored population and datasets

The study was carried out on the king penguin *Aptenodytes patagonicus* colony of ‘Baie du Marin’ settled on the Possession Island in the Crozet Archipelago (46°25S, 51°45E). A sub-area of the colony of ca. 10,000 breeding pairs has been continuously electronically monitored since 1998 using Radio-Frequency IDentification (RFID) technology (Bardon et al., 2023; Gendner et al., 2005). As of 2024, the database contains information on ∼17,000 birds, including ∼12,000 of known age and ∼5,500 currently active in the colony.

Using a new application of deep learning (Bardon et al., 2023) on our RFID-detection dataset, we estimated the status and the timing of breeding for each micro-tagged individual and each breeding season. Briefly, we used a convolutional neural network workflow to classify each season based on a 10-year training of direct field observations and performed two testing steps to ensure the quality of the classification (see Bardon et al. (2023) for precision on the system and on classification methods). For each individual and each year, we thus obtained the breeding status (non-breeding, failed breeding, successful breeding) and the most probable date for the onset of breeding (defined here as the mating event). King penguins breed in an asynchronous cycle that lasts from 12 to 14 months (Barrat, 1976). As a result, every year, the successful individuals of the previous year either take a “sabbatical” or start breeding later in the season (Jiguet & Jouventin, 1999).To avoid confounding our results with the variation in asynchrony parameters, we restricted our dataset to birds that (1) did not breed successfully in the previous season, (2) were at least 9 years old at the start of the breeding season, since young inexperienced breeders have a low overall contribution to the breeding success of the population (see Supp. Figure 2 and Supp. Figure 3). We included in this dataset unknown-age adults, as these birds were systematically RFID-tagged at an advanced stage of breeding, they were at least 5 years old (the mean age at first breeding of sexual maturity), and we then used data of these birds with a 3-year delay after their marking. Out of the 28,000 breeding seasons available in our dataset, 8,800 from around 2,600 individuals were thus selected to compute the annual median breeding initiation date of the colony and the annual breeding success rate, obtained by dividing the number of successful cycles by the number of breeding attempts (Supp. Figure 4).

### Environmental datasets

We selected climate variables likely to have an impact on the phenology and reproductive success of king penguins from the Crozet population (see Supp. Table 1). Two main areas were considered: 1) an area centred on Crozet archipelago (43°S-47°S – 46°E-56°E) used by the birds during their foraging circular trips (sensu Jouventin et al., 1994) that we refer to as “Crozet Area”; 2) an area south of the archipelago (47°S-53°S – 49°E-55°E) corresponding to their main foraging area during summer, tightly linked to the polar front (Bost et al., 2015; Charrassin & Bost, 2001), that we refer that as “Polar Front Area”.

Sea surface temperature (SST) is widely reported as one of the main drivers of marine productivity, trophic interactions, and distribution of species (Cheung et al., 2013). SST is likely to impact breeding phenology and success as it has been already shown previously for many seabirds (Carroll et al., 2015) and in particular for several penguin species (Cullen et al., 2009; Cunningham & Moors, 1994; Le Bohec et al., 2008). We computed daily mean OI (optimal interpolation) SST in the areas around and south of the archipelago. We computed 2°, 4°, and 5° isotherm latitude south of Crozet as they were also previously linked to the positions of the Antarctic polar fronts and the main foraging area of the king penguins (Bost et al., 2015; Park et al., 1993). We considered the distance between isotherms 4° and 5° as an index of SST latitudinal gradient between the breeding colony and the main foraging area. The latitude of the Antarctic Polar Front has been defined as the northernmost extent of the 200 m surface layer temperature minimum of 2 °C (Pollard et al., 2002) and computed at the latitude of Crozet from the GODAS model (NCEP Global Ocean Data Assimilation System). The latitude of the northern edge of the Marginal Ice Zone (MIZ), defined as the area covered by an ice concentration between 15 and 80%, was considered as previous studies showed presence of king penguins at these latitudes, especially during the winter season (Bost et al., 2004; Charrassin & Bost, 2001). In addition to the MIZ latitude, we computed the part of the total ice area (with ice concentration >15%) that is covered by the marginal ice zone (between 15% and 80% of sea ice concentration) in the defined area south of Crozet (50°S-70°S - 46°E-56°E), as an index of the quality of the sea ice to forage.

As a proxy of marine productivity, we considered the mean concentration of Chlorophyll *a* (Chl*a*) both in Crozet Area and in the Polar Front Area, which could act as a phenological cue (Crossin et al., 2022) and index of prey availability. We computed the centre of mass of the Chl*a* concentrations at the longitude of Crozet and between 40°S and 56°S as an indicator of location of the maximum productivity.

Wind force and storm probability were used to reflect direct climatic constraints on land and at sea that can affect breeding birds and their chicks. Wind force index was computed from the daily maximum wind force by taking the monthly mean of the 25% windiest days. The storm track probability index was computed as the mean latitude of the maximum gradient in the isobaric surface 500 mb (Marshall & Plumb, 2007).

Finally, two global climatic indices were considered to detect indirect effects of climate that can be left unexplained by more specific climate variables. The Southern Annular Mode (SAM) defined as 700mb height anomalies poleward of 20°S onto the loading pattern of the Antarctic Oscillation (the leading mode of Empirical Orthogonal Function (EOF) analysis of monthly mean 700 hPa height during 1979-2000 period) was considered as the main mode of this region (D. W. J. Thompson & Wallace, 2000). In addition, the Southern Oscillation Index (SOI), which reflects large-scale fluctuations in air pressure between Tahiti and Darwin and serves as a proxy for El Niño–Southern Oscillation (ENSO) phases, was used to account for potential remote climatic influences from the tropical Pacific region.

### General trends

Phenological annual trend, given by the annual median breeding date, has been tested using a linear and a quadratic regression of the phenology against the year using lm() function in R (version 4.3.0 R Core Team, 2023). We considered that the quadratic regression best supported the correlation if the Akaike information criterion corrected for small sample size (AICc) were at least inferior to the linear regression by 2 units (Akaike, 1998; Anderson & Burnham, 2004).

Trend of the mean annual breeding success rate was tested using a linear regression of the mean annual breeding success against the year, treated as a quantitative variable. As breeding success is likely linked to the breeding starting date, we also tested if the temporal trend of the annual breeding success rate is explained by the annual median breeding date. Although reproductive success is a proportion theoretically bounded between 0 and 1, the observed values in our dataset ranged between 0.3 and 0.7. Within this restricted interval, we considered linear and quadratic regression models to be appropriate approximations for investigating the relationships between population reproductive success and environmental conditions.

### Time window identification

Despite previous studies relating environmental predictors with annual breeding traits of penguins, it is still unclear when these climate variables can affect breeding. To avoid arbitrary choice of temporal windows and explore possible long-term effects, we used a systematic method to find which time window has the strongest association with the response variable, while avoiding false-positive correlations when testing many variables. We therefore used the sliding window approach implemented in R (version 4.3.0, R Core Team, 2013) in the *Climwin* package (Bailey & Van De Pol, 2016; Van De Pol et al., 2016).

We first defined baseline models for phenology and breeding success. To avoid spurious correlation due to temporal trends (Iler et al., 2017), we included year as a continuous variable in both baseline models. For breeding success, we included the median breeding date in the baseline model as it greatly explained the annual breeding success (R² = 0.60). For both phenology and breeding success, we compared the AICc given by the linear and quadratic model with different time windows of each climate variable with the respective baseline models. All combinations of time windows in the temporal extent that we considered were tested. For phenology, we considered a period of 30 weeks before the breeding season (1^st^ of November), a period long enough to catch short-lagged effects of climate conditions while avoiding autocorrelation effects between seasons.

For breeding success, because we designed our dataset to remove the breeding birds that were successful the previous season, there was no correlation between the current and the past breeding success rate. We then considered a longer period of 2 years preceding the beginning of the breeding season and until 1 year after (corresponding to the end of a successful breeding season).

The best model was chosen for each response variable and each environmental explanatory variable based on the highest difference in AICc (ΔAICc) with the baseline models (Akaike, 1998; Anderson & Burnham, 2004). For each environmental variable, the quadratic model was selected only when the ΔAICc was at least 2 units higher than the best linear model, otherwise the linear model was selected.

To control for multiple testing across numerous time windows, we used Climwin’s randomization procedure (Van De Pol et al., 2016). This approach shuffles the response variable 10 times while maintaining the temporal structure of environmental predictors, generating a null distribution to estimate the probability (Pc) that observed correlations could arise by chance alone. Only variables with Pc < 0.5 were retained, reducing the risk of false positives inherent in sliding window analyses.

To reduce the risk of overfitting arising from a high number of candidate variables relative to the number of data points, we first assessed multicollinearity among variables using variance inflation factors (VIF). The variables with the highest VIF were removed iteratively until no variables left had a VIF > 3. Then, we performed a model selection with all the candidate variables using the package *MuMin* and *Dredge* function (Bartoń, 2023). Among all equally best performing models (with a ΔAICc < 2 from the lowest AICc model), we selected the one with the lowest number of parameters (Anderson & Burnham, 2004).

Partial regression plots are given for the best models. They are constructed by plotting the residuals of the dependent variable (after removing the effects of all other predictors) against the residuals of the independent variable of interest (after similarly controlling for other predictors). We then added the mean value of the dependent variable and of the variable of interest.

### Contribution of environmental conditions in explaining observed trends

We estimated how much of the observed trends could be explained by the environmental variables obtained from our model selection. We added the year variable in the final selected model and extracted its remaining associated linear coefficient. We compared this coefficient to the slope of the temporal trend-only model to obtain the part of the observed temporal trend explained in the final model including environmental variables (Haest et al., 2020).

## Acknowledgments

This study was supported by the Institut Polaire Français Paul-Emile Victor (IPEV) within the framework of the project 137-ANTAVIA, by the Centre Scientifique de Monaco, and by the Centre National de la Recherche Scientifique (CNRS) through the Programme Zone Atelier de Recherches sur l’Environnement Antarctique et Terres Australes Subantarctique (ZATA), by Research Council of Finland (#331320 and #354649), and by the Canada Research Program, NSERC, and Université de Moncton. This study is part of and supported by the long-term Studies in Ecology and Evolution (SEE-Life) programme of CNRS. We are deeply grateful to all the wintering and summering members of project IPEV 137 and all the other colleagues and students within the team, who participated in the long-term monitoring as part of the project IPEV 137 since 1998. All members of the MIBE team at the IPHC in Strasbourg are thanked for their technical expertise and support. We also sincerely thank the IPEV logistics teams for their important and continued support in the field. This study was approved by the French Polar Environmental Committee and permits handling animals and access breeding sites were delivered by the Terres Australes et Antarctiques Françaises (TAAF).

## Author contributions

Conceptualization: G.B., R.C., C.L.B. Methodology: G.B., J.M.D., R.C., C.L.B. Data analysis: G.B., R.C. Data collection: G.B., N.C., Y.L.M., R.C., C.L.B.. Supervision: R.C., C.L.B.. Writing—original draft: G.B., R.C., C.L.B. All authors contributed critically to the drafts and gave final approval for publication.

## Ethics

Health, safety, security and other risks for participating researchers were assessed and managed by the institution (Institut Polaire Français Paul-Emile Victor, IPEV) that provided logistic support at the research stations involved. Ethics questions concerning local and regional researchers, partners, or governments do not apply, due to Sub-Antarctic Islands being uninhabited.

