## Supplementary Information for "Multiannual environmental forcing shapes breeding phenology and success in a subantarctic seabird"

* Co-last authors by alphabetical order

### Supplementary Information

#### Supplement A. Study area

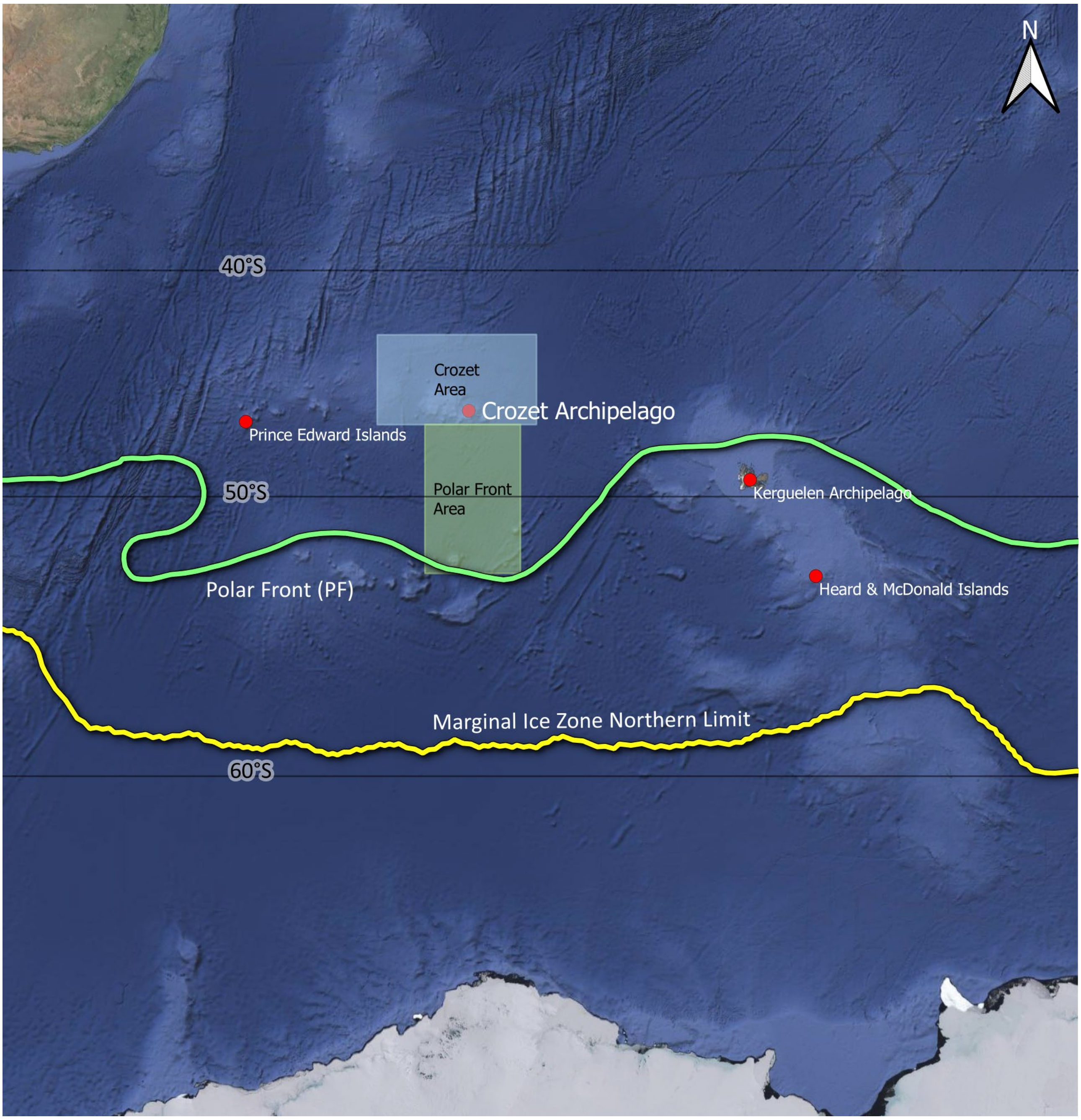


*Supp. Figure 1.* ***Location of the study site (Crozet Archipelago) in the Southern Ocean.*** *Other islands in this area with king penguin colonies are also illustrated. The Crozet Area (43S-47S - 46E-56E) and the Polar Front Area (47S-53S - 49E-55E) are the areas used to calculate our environmental variables (Sea Surface Temperature SST, and Chlorophyll a concentrations Chla). The location of the polar front and the northern limit of the Marginal Ice Zone (MIZ) were extracted from the Quantarctica datasets (Matsuoka et al., 2021). The northern limit of the MIZ was calculated from the mean sea ice extent in October between 1981 and 2010 (Fetterer et al., 2017). The polar front location represents the climatological position and was estimated from observed temperature and salinity data (Orsi et al., 1995). The background map was extracted from GoogleEarth.*

#### Supplement B. Dataset selection

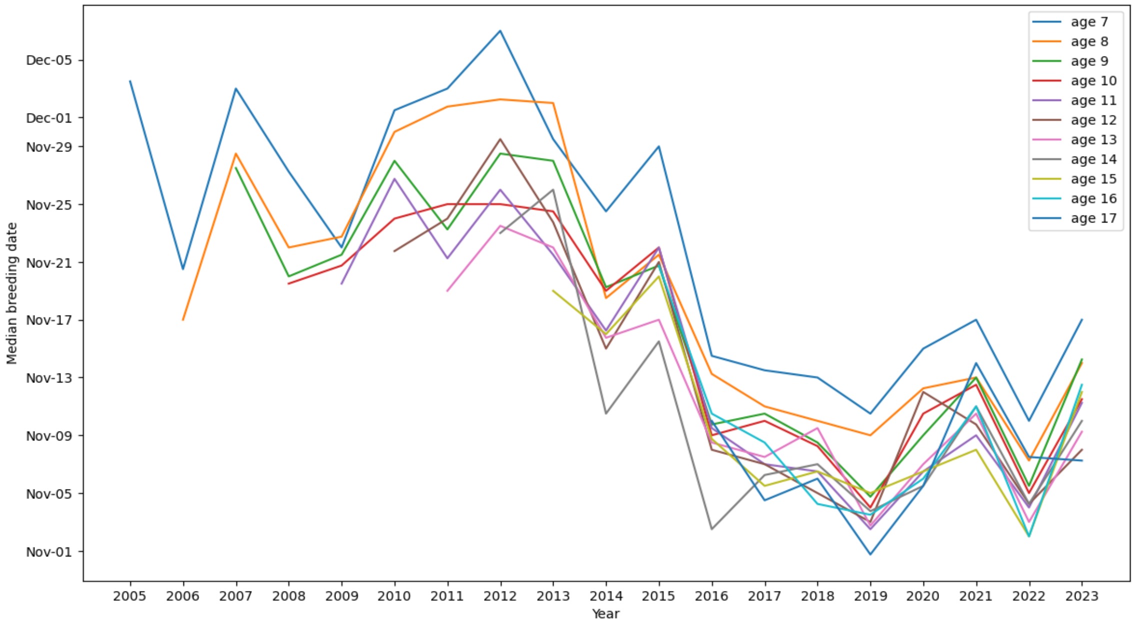


*Supp. Figure 2.* ***Median breeding initiation date calculated for groups of king penguins of any breeding ages and for each monitoring year.*** *Birds aged 7 and 8 (the first effective breeding ages for king penguins) breed later each year than older groups and have therefore been removed from the dataset used in the study.*

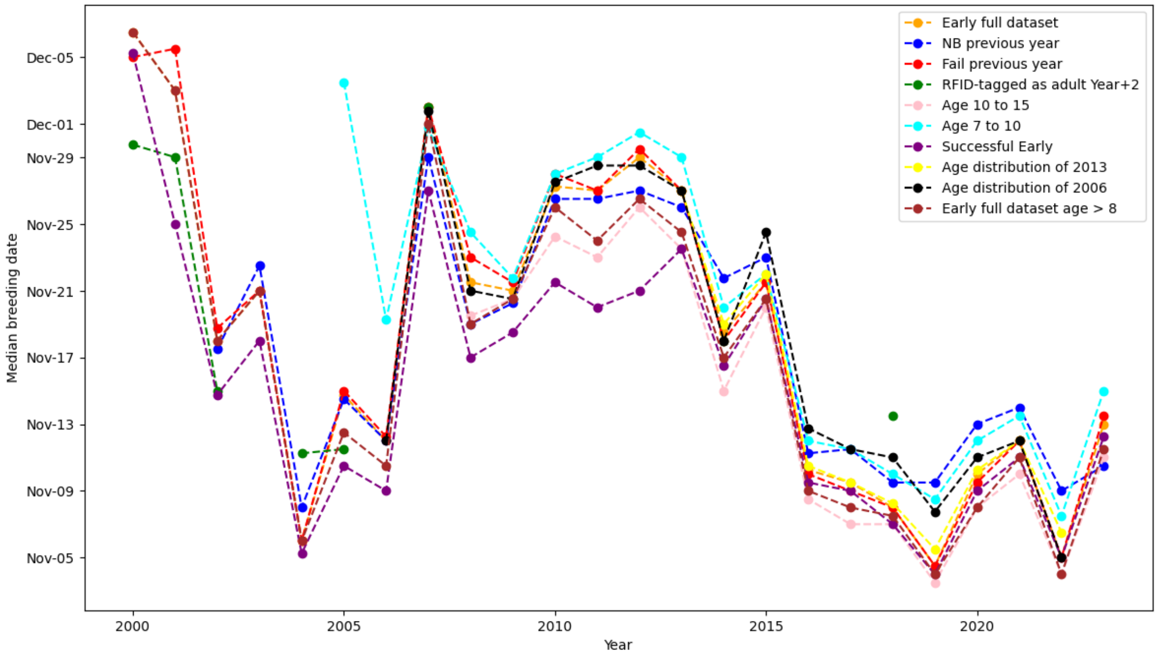


*Supp. Figure 3.* ***Median breeding initiation date calculated for groups of king penguins with various filters****. 'Early full dataset': all birds considered 'early' (not successful the previous year). 'NB previous year': all birds that were non-breeder the previous year. 'Fail previous year': all birds that failed to breed the previous year. 'RFID-tagged as adult Year N+2': only early birds that were tagged as adults (with unknown age) and 2 years after their tagging (because we do not have the breeding starting date of year N and most of them are not 'Early' during year N+1). 'Age 10 to 15': only early birds aged between 10 and 15. 'Age 7 to 10': only early birds aged between 7 and 10. 'Successful Early': birds that were successful in year N. 'Age distribution of 2013': resampling of birds to keep the known age distribution of 2013. 'Age distribution of 2006': resampling of birds to keep the known age distribution of 2006. 'Early full dataset age > 8': dataset used in the study with only early birds aged more than 8.*

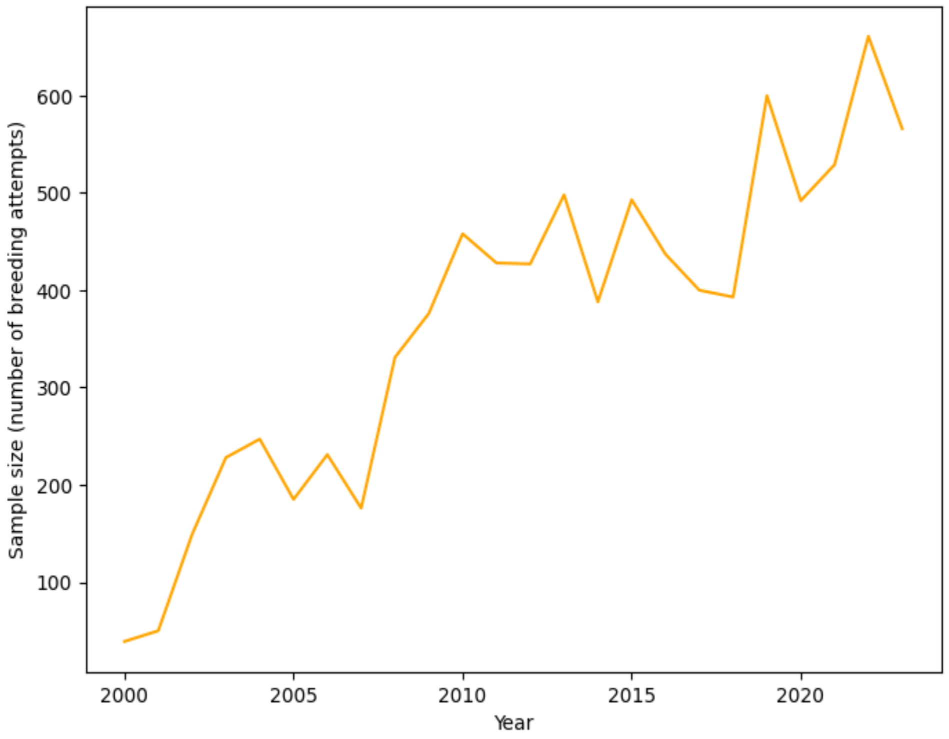


*Supp. Figure 4.* ***Number of breeding attempts per year in the final dataset containing king penguins aged more than 8 and being 'early'.***

#### Supplement C. Climate variables

*Supp. Table 1.* ***Summary of climate variables tested in the study.***

| Variable | Extent | Time resolution | Data source |
|---|---|---|---|
| Optimal Interpolation Sea Surface Temperature (OISST) in the Crozet Area | 43°S-47°S - 46°E-56°E | Daily | https://www.ncei.noaa.gov/products/optimum-interpolation-sst |
| OISST in the Polar Front Area | 47°S-53°S - 49°E-55°E | Daily | https://www.ncei.noaa.gov/products/optimum-interpolation-sst |
| Isotherm 2° SST latitude | South at the longitude of Crozet (51°E) | Daily | https://www.ncei.noaa.gov/products/optimum-interpolation-sst |
| Isotherm 4° SST latitude | South at the longitude of Crozet (51°E) | Daily | https://www.ncei.noaa.gov/products/optimum-interpolation-sst |
| Isotherm 5° SST latitude | South at the longitude of Crozet (51°E) | Daily | https://www.ncei.noaa.gov/products/optimum-interpolation-sst |
| Distance between isotherms 4° and 5°SST (SST gradient) | South at the longitude of Crozet (51°E) | Daily | https://www.ncei.noaa.gov/products/optimum-interpolation-sst |
| Southern Annular Mode (SAM) | Global | Daily | https://psl.noaa.gov/data/20thC_Rean/timeseries/monthly/SAM/ |
| Marginal Ice Zone (MIZ) latitude | 46°E-56°E | Daily | https://noaadata.apps.nsidc.org/NOAA/G02202_V4/south/aggregate/ |
| Mean wind force around Crozet | 44°S - 48°S - 50°E - 54°E | Monthly | https://www.psl.noaa.gov/data/gridded/data.ncep.reanalysis.html |
| Index of storm probability | 30°S to 75°S - At the longitude of Crozet (51°E) | Monthly | https://www.psl.noaa.gov/data/gridded/data.ncep.reanalysis.html |
| Antarctic Polar Front latitude (limit of 2°C water in the upper watermass) | South at the longitude of Crozet (51°E) | Monthly | https://www.psl.noaa.gov/data/gridded/data.godas.html |
| Mean Chlorophyll *a* concentration (Chl*a*) in the Crozet Area | 43°S-47°S - 46°E-56°E | Daily | Global Ocean Colour (Copernicus-GlobColour), Bio-Geo-Chemical, L4 (monthly and interpolated) from Satellite Observations |
| Mean Chl*a* in the Polar Front Area | 47°S-53°S - 49°E-55°E | Daily | Global Ocean Colour (Copernicus-GlobColour), Bio-Geo-Chemical, L4 (monthly and interpolated) from Satellite Observations |
| Centre of mass of Chl*a* in the Southern Ocean | 40°S to 54°S - At the longitude of Crozet (51°E) | Daily | Global Ocean Colour (Copernicus-GlobColour), Bio-Geo-Chemical, L4 (monthly and interpolated) from Satellite Observations |
| Southern Oscillation Index (SOI) | Global | Daily | https://data.longpaddock.qld.gov.au/SeasonalClimateOutlook/SouthernOscillationIndex/SOIDataFiles/DailySOI1933-1992Base.csv |

#### Supplement D. Model selection

**Summary of climate variables tested in the study for phenology analysis.**

The first step of variable selection resulted in candidate time windows for nine variables. From those, four were removed due to correlation with other variables. We then conducted the model selection with the following variables: the mean Sea Surface Temperature (SST) in the Polar Front Area during autumn, the mean SST in the Crozet Area during winter, the Marginal Ice Zone (MIZ) latitude during autumn and winter, the SOI during autumn, and the mean Chlorophyll *a* concentration in the Polar Front Area at the end of winter (see Supp. Table 2).

*Supp. Table 2.* ***Model selection for prediction of king penguin phenology.*** *Chla PFA: Chlorophyll a Polar Front Area, SST PFA: Sea Surface Temperature Polar Front Area, SOI: Southern Oscillation Index, MIZ lat: Marginal Ice Zone Latitude.*

| Covariate | df | AICc | ΔAICc | Weight |
|---|---|---|---|---|
| Chla PFA + SST PFA² | 5 | 146.4 | 0 | 0.718 |
| Chla PFA + SST PFA² + SOI | 6 | 149.3 | 2.96 | 0.163 |
| Chla PFA + SST PFA² + MIZ Lat | 6 | 149.9 | 3.59 | 0.119 |
| Year | 3 | 170.9 | 24.5 | |

**Summary of climate variables tested in the study for breeding success analysis.**

The first step of variable selection resulted in candidate climate windows for 7 variables (see Supp. Table 3). From those, two were removed due to correlation with another variable. We then conducted the model selection with the following variables: mean Chlorophyll *a* concentration in the Polar Front Area, mean Chlorophyll *a* concentration in the Crozet Area, latitude of the 5° isotherm, distance between isotherms 4° and 5°, Southern Annular Mode (SAM), Index of storm track probability (see Supp. Table 3).

*Supp. Table 3.* ***Model selection for prediction of king penguin breeding success.*** *Phenology: Annual median breeding date, SAM: Southern Annular Mode index, SST Gradient: Distance between 4°C and 5°C isotherms, Chla Crozet Area: Chlorophyll a concentration in Crozet Area, Storm Index: Storm track probability index, 5°C Lat: Latitude of 5°C isotherm, Chla Polar Front Area: Chlorophyll a concentration in the Polar Front Area.*

| Covariate | df | AICc | ΔAICc | Weight |
|---|---|---|---|---|
| Phenology + SAM + SST Gradient + CHla Crozet Area² | 7 | -82.1 | 0 | 0.590 |
| Phenology + SST Gradient + CHla Crozet Area² | 6 | -79.9 | 2.16 | 0.201 |
| Phenology + SAM + SST Gradient + CHla Crozet Area² + 5°C Lat | 8 | -78.7 | 3.38 | 0.109 |
| Phenology + SST Gradient + CHla Crozet Area² + CHla Polar Front Area² | 8 | -78.6 | 3.54 | 0.101 |

#### Supplement E. Supplement results for phenology analysis

*Supp. Table 4.* ***Final selected model summary for king penguin phenology.*** *Multiple linear regression of the annual median breeding date according to the chlorophyll concentration in the Polar Front Area (Chla_PFA) and to the Sea Surface Temperature in the Polar Front Area (SST_PFA). Multiple R-squared: 0.8243, Adjusted R-squared: 0.798*

| | Estimate | Std. Error | Pr(>\|t\|) |
|---|---|---|---|
| Intercept | 953.734 | 160.950 | 8.53e-06 *** |
| Chla_PFA | 571.155 | 111.643 | 5.27e-05 *** |
| SST_PFA | -449.287 | 71.250 | 3.72e-06 *** |
| $SST\_PFA^2$ | 49.598 | 8.024 | 4.88e-06 *** |

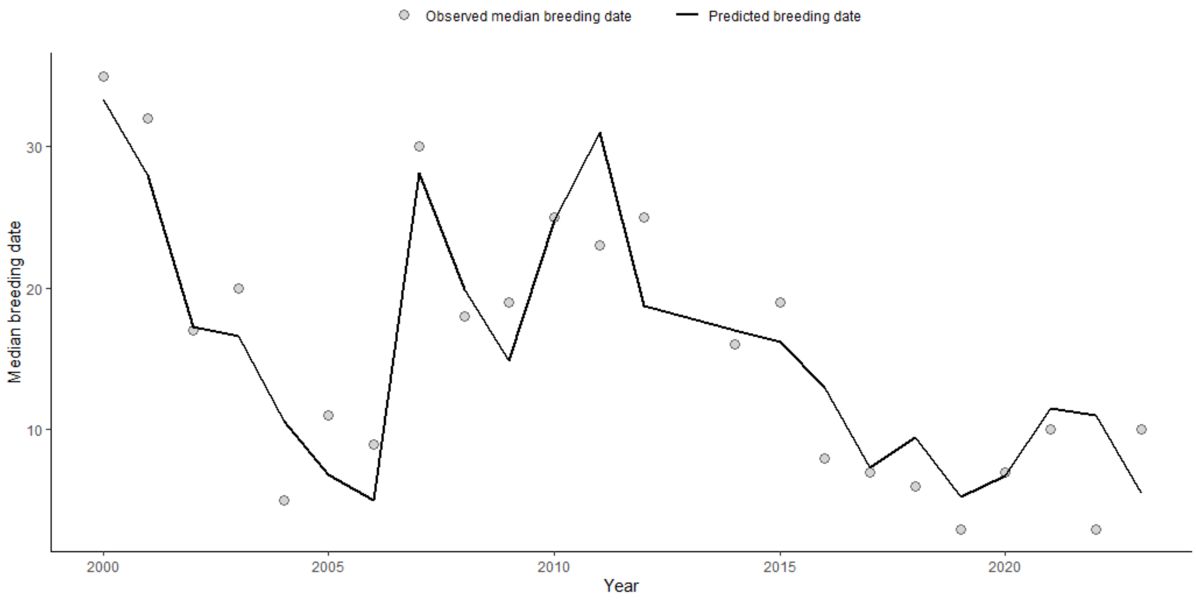


*Supp. Figure 5.* ***Observed and predicted breeding initiation date in king penguins, Crozet Archipelago.***

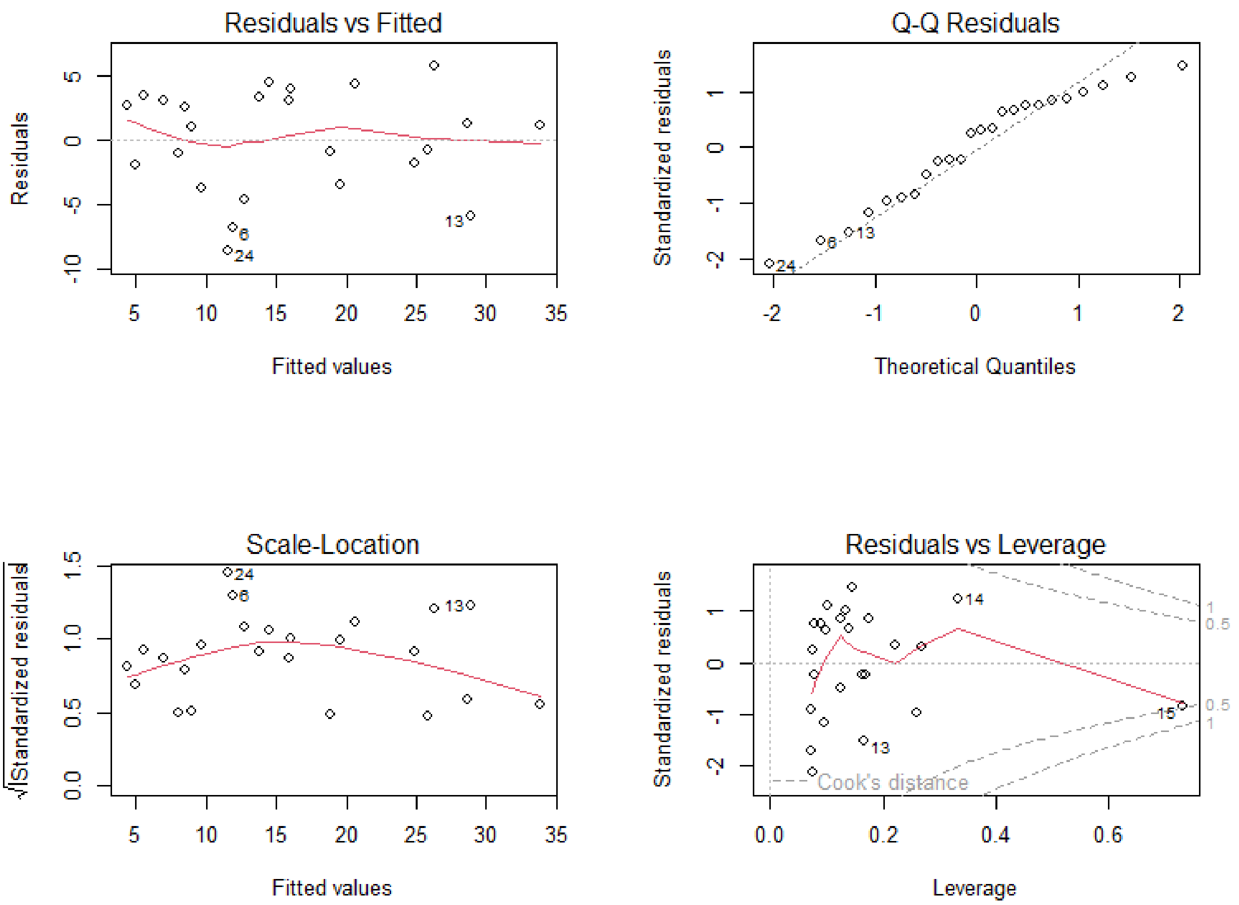


*Supp. Figure 6.* ***Diagnostic plots of the best model for king penguin phenology analysis.*** *(A) Residuals vs Fitted; (B) Normal Q-Q; (C) Scale-Location; (D) Cook's Distance. These plots indicate that model assumptions were reasonably met.*

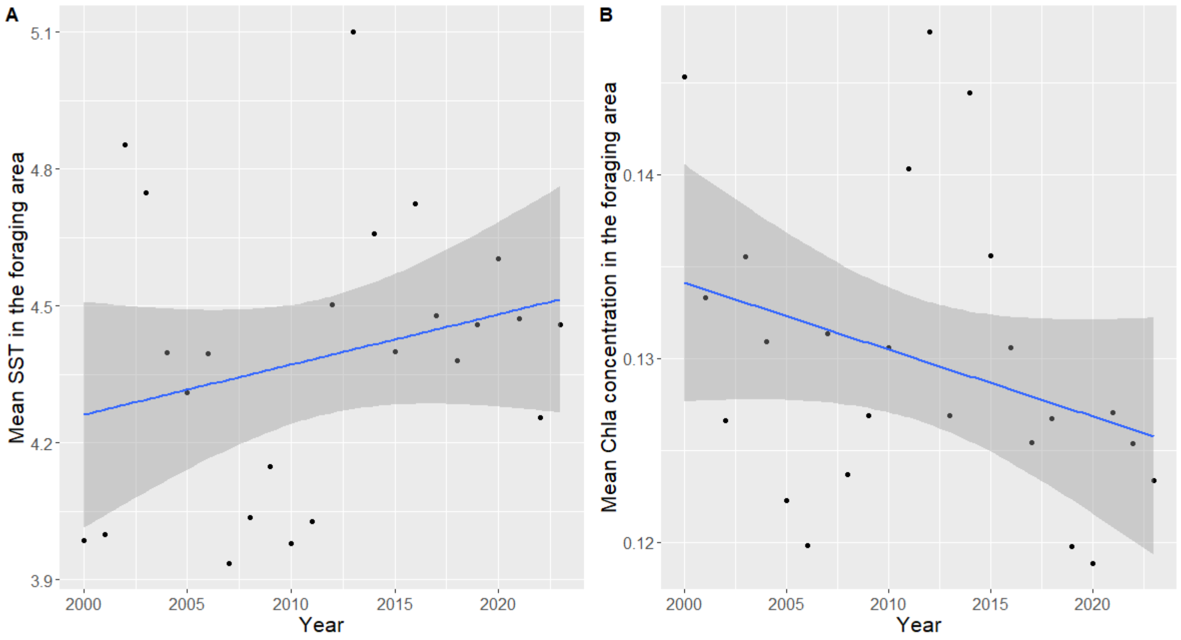


*Supp. Figure 7. **Temporal trends of the best selected environmental variables explaining breeding phenology in king penguins, Crozet Archipelago.** (A) represents the trend for the mean Sea Surface Temperature (SST) in the Polar Front Area and during the best temporal window, i.e. in the fall preceding the breeding season. (B) represents the trend for the mean Chlorophyll a concentration (Chla) in the Polar Front Area and during the best temporal window, i.e. in the spring preceding the breeding season. Each point represents one year and the blue line simple linear regressions. The grey shaded areas show 95% confidence intervals of the trend lines. P-values of year coefficient are 0.217 and 0.1313 for (A) and (B), respectively. Slopes of the year coefficient are 0.011 ± 0.0089 (p-value = 0.217) and -0.00036 ± 0.00023 (p-value = 0.1313) for (A) and (B) respectively.*

#### Supplement F. Supplement results for breeding success analysis

*Supp. Table 5.* ***Final selected model summary for king penguin phenology.*** *Multiple linear regression of the annual breeding success rate according to the annual median breeding date (Phenology), the gradient of Sea Surface Temperature south of Crozet (SST_Gradient), the Southern Annular Mode index (SAM) and the Chlorophyll a concentration in the Crozet Area (Chla_Crozet_Area). Multiple R-squared: 0.9391, Adjusted R-squared: 0.9221*

| | Estimate | Std. Error | Pr(>\|t\|) |
|---|---|---|---|
| Intercept | 0.8379550 | 0.0544418 | 8.37e-12 *** |
| Phenology | -0.0103023 | 0.0007818 | 1.10e-10 *** |
| SST_Gradient | -0.0862725 | 0.0244353 | 0.00239 ** |
| SAM | 0.0339748 | 0.0146494 | 0.03235 * |
| CHla_Crozet_Area | 0.0967712 | 0.0353152 | 0.01345 * |
| $CHla\_Crozet\_Area^2$ | -0.1307176 | 0.0425171 | 0.00653 ** |

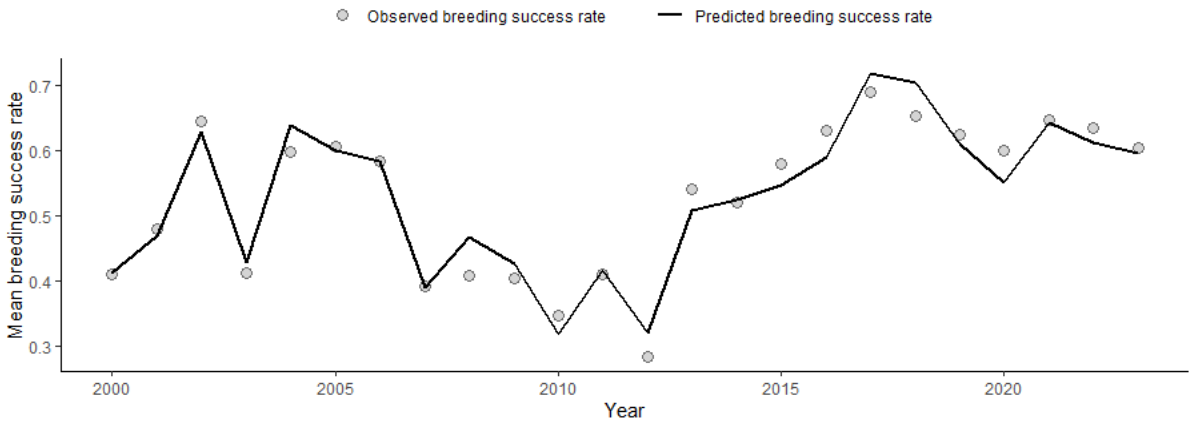


*Supp. Figure 8.* ***Observed and predicted breeding success rate in king penguins, Crozet Archipelago.***

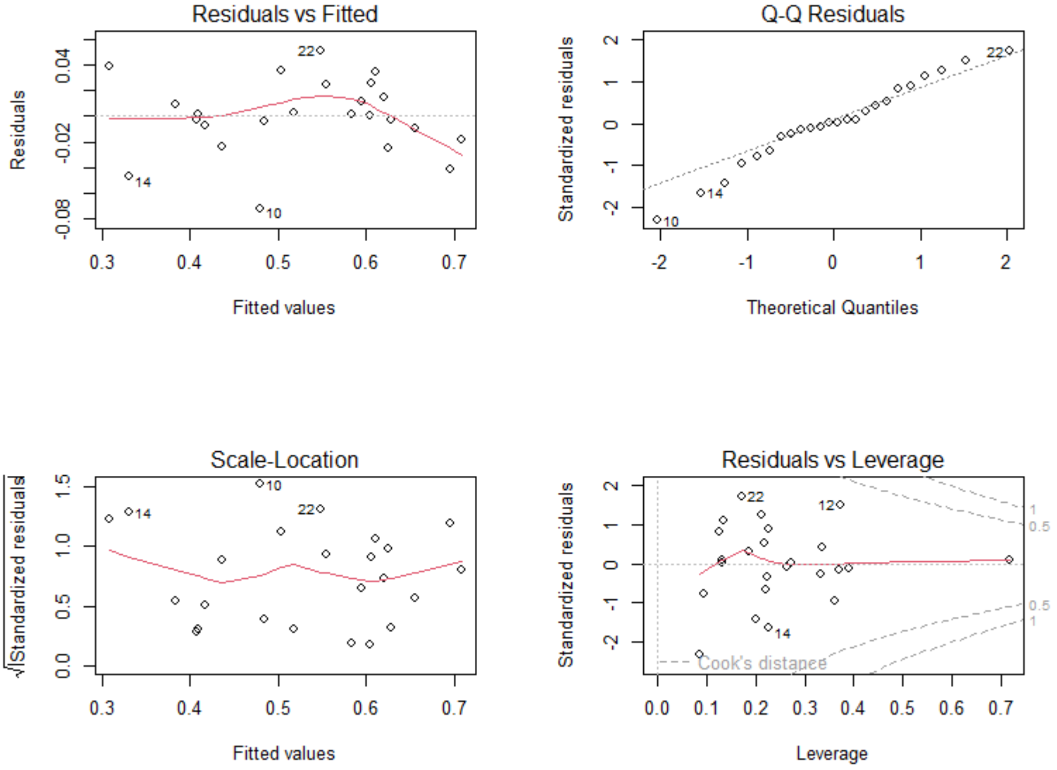


*Supp. Figure 9.* ***Diagnostic plots of the best model for king penguin breeding success analysis.*** *(A) Residuals vs Fitted; (B) Normal Q-Q; (C) Scale-Location; (D) Cook's Distance. These plots indicate that model assumptions were reasonably met.*

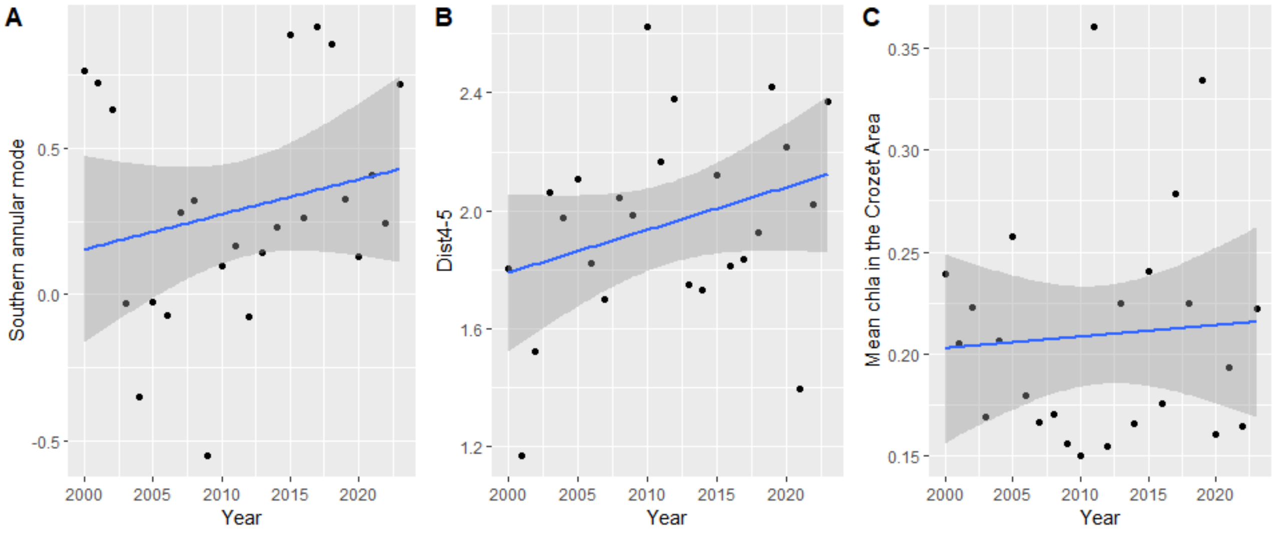


*Supp. Figure 10.* ***Temporal trends of the best selected environmental variables explaining breeding success in king penguins, Crozet Archipelago.*** *(A) represents the trend for the mean Sea Surface Temperature (SST) in the Southern Annular Mode index (SAM) and during the best temporal window, i.e. from February to May (B) represents the trend for the distance between the isotherms 4°C and 5°C during the best temporal window, i.e. from mid-December to mid-April, (C) mean Chlorophyll a concentration in the Crozet Area and during the best temporal window, i.e. from mid-August to early September. Each point represents one year and the blue line simple linear regressions. The grey shaded areas show 95% confidence intervals of the regression trend lines. Slopes of the year coefficient are 0.012 ± 0.011 (p-value = 0.31), 0.0145 ± 0.0095 (p-value = 0.143) and 0.00056 ± 0.00167 (p-value = 0.74) for (A), (B) and (C) respectively.*

#### Supplement G. Variable selection for phenology

*Supp. Table 6.* ***Summary of the variable selection and time window identification using Climwin for king penguin phenology.*** *Highlighted rows correspond to the candidate variable with Pc (the probability that a signal is not due to chance) < 0.5. The quadratic model was selected only when the ΔAICc was at least 2 units higher than the best linear model, otherwise the linear model was selected.*

| Tested variable | Regression type | ΔAICc | Pc |
|---|---|---|---|
| SST PFA | Linear | 2.06 | 0.78 |
| SST PFA | Quadratic | 9.37 | 0.30 |
| SST Cro | Linear | 3.52 | 0.65 |
| SST Cro | Quadratic | 8.83 | 0.11 |
| 2°C Iso Lat | Linear | 3.07 | 0.72 |
| 2°C Isotherm Lat | Quadratic | 10.14 | 0.10 |
| 4°C Isotherm Lat | Linear | 1.06 | 0.75 |
| 4°C Isotherm Lat | Quadratic | 8.56 | 0.08 |
| 5°C Isotherm Lat | Linear | 3.23 | 0.61 |
| 5°C Isotherm Lat | Quadratic | 7.99 | 0.37 |
| Chla PFA | Linear | 7.14 | 0.42 |
| Chla_PFA | Quadratic | 5.39 | 0.71 |
| Chla Cro | Linear | -1.76 | 0.81 |
| Chla Cro | Quadratic | -1.98 | 0.86 |
| Chla Mass Centre | Linear | 4.61 | 0.55 |
| Chla Mass Centre | Quadratic | 6.34 | 0.50 |
| SST Gradient | Linear | 9.27 | 0.28 |
| SST Gradient | Quadratic | 6.35 | 0.50 |
| SAM | Linear | 3.25 | 0.74 |
| SAM | Quadratic | 3.12 | 0.77 |
| MIZ Lat | Linear | 5.83 | 0.25 |
| MIZ Lat | Quadratic | 3.34 | 0.52 |
| Wind | Linear | 1.13 | 0.76 |
| Wind | Quadratic | -1.04 | 0.91 |
| Storm | Linear | 3.75 | 0.73 |
| Storm | Quadratic | 0.66 | 0.90 |
| APF Lat | Linear | -1.01 | 0.82 |
| APF Lat | Quadratic | -3.74 | 0.86 |
| SOI | Linear | 18.32 | 0.007 |
| SOI | Quadratic | 15.36 | 0.003 |

#### Supplement F. Variable selection for breeding success

*Supp. Table 7.* ***Summary of the variable selection and time window identification using Climwin for king penguin breeding success.***

| Tested variable | Regression type | ΔAICc | Pc |
|---|---|---|---|
| SST PFA | Linear | 6.39 | 0.78 |
| SST PFA | Quadratic | 9.8 | 0.91 |
| SST Cro | Linear | 5.91 | 0.58 |
| SST Cro | Quadratic | 11.25 | 0.88 |
| 2°C Iso Lat | Linear | 5.65 | 0.85 |
| 2°C Isotherm Lat | Quadratic | 7.68 | 0.66 |
| 4°C Isotherm Lat | Linear | 8.5 | 0.77 |
| 4°C Isotherm Lat | Quadratic | 7.54 | 0.92 |
| 5°C Isotherm Lat | Linear | 8.45 | 0.38 |
| 5°C Isotherm Lat | Quadratic | 15.97 | 0.27 |
| Chla PFA | Linear | 3.56 | 0.76 |
| Chla_PFA | Quadratic | 9.65 | 0.05 |
| Chla Cro | Linear | 11.74 | 0.10 |
| Chla Cro | Quadratic | 18.49 | 0.29 |
| Chla Mass Centre | Linear | 3.59 | 0.83 |
| Chla Mass Centre | Quadratic | 6.06 | 0.99 |
| SST Gradient | Linear | 11.74 | 0.04 |
| SST Gradient | Quadratic | 9.75 | 0.29 |
| SAM | Linear | 14.87 | 0.08 |
| SAM | Quadratic | 11.26 | 0.08 |
| MIZ Lat | Linear | 6.76 | 0.47 |
| MIZ Lat | Quadratic | 11.41 | 0.05 |
| Wind | Linear | 3.25 | 0.93 |
| Wind | Quadratic | 0.02 | 0.94 |
| Storm | Linear | 11.86 | 0.51 |
| Storm | Quadratic | 12.41 | 0.075 |
| APF Lat | Linear | -1.15 | 0.82 |
| APF Lat | Quadratic | 3.34 | 0.73 |
| SOI | Linear | 3.66 | 0.82 |
| SOI | Quadratic | 5.9 | 0.91 |

#### Supplementary information references


Fetterer, F., Knowles, K., Meier, W., Savoie, M., & Windnagel, A. (2017). *Sea Ice Index, Version 3* [Dataset]. NSIDC. https://doi.org/10.7265/N5K072F8

Matsuoka, K., Skoglund, A., Roth, G., de Pomereu, J., Griffiths, H., Headland, R., Herried, B., Katsumata, K., Le Brocq, A., Licht, K., Morgan, F., Neff, P. D., Ritz, C., Scheinert, M., Tamura, T., Van de Putte, A., van den Broeke, M., von Deschwanden, A., Deschamps-Berger, C., … Melvær, Y. (2021). Quantarctica, an integrated mapping environment for Antarctica, the Southern Ocean, and sub-Antarctic islands. *Environmental Modelling & Software*, *140*, 105015. https://doi.org/10.1016/j.envsoft.2021.105015

Orsi, A. H., Whitworth, T., & Nowlin, W. D. (1995). On the meridional extent and fronts of the Antarctic Circumpolar Current. *Deep Sea Research Part I: Oceanographic Research Papers*, *42*(5), 641–673. https://doi.org/10.1016/0967-0637(95)00021-W
